# Retinal waves in adaptive rewiring networks orchestrate convergence and divergence in the visual system

**DOI:** 10.1101/2023.10.11.561834

**Authors:** Raul Luna, Jia Li, Roman Bauer, Cees van Leeuwen

## Abstract

Spontaneous retinal wave activity shaping the visual system is a complex neurodevelopmental phenomenon. Retinal ganglion cells are the hubs through which activity diverges throughout the visual system. We consider how these divergent hubs emerge, using an adaptively rewiring neural network model. Adaptive rewiring models show in a principled way how brains could achieve their complex topologies. Modular small-world structures with rich club effects and circuits of convergent-divergent units emerge as networks evolve, driven by their own spontaneous activity. Arbitrary nodes of an initially random model network were designated as retinal ganglion cells. They were intermittently exposed to the retinal waveform, as the network evolved through adaptive rewiring. A significant proportion of these nodes developed into divergent hubs within the characteristic complex network architecture. The proportion depends parametrically on the wave incidence rate. Higher rates increase the likelihood of hub formation, while increasing the potential of ganglion cell death. In addition, direct neigbours of designated ganglion cells differentiate like amacrine cells. The divergence observed in ganglion cells resulted in enhanced convergence downstream, suggesting that retinal waves control the formation of convergence in LGN. We conclude that retinal waves stochastically control the distribution of converging and diverging activity in evolving complex networks.

## Introduction

The connectivity of the central nervous system is a work in progress. During development and learning, synapses are being added, removed, and remodeled in response to neural network activity. Although later changes are thought to be largely experience-dependent (e.g. Himmelberg et al., 2023), a prominent role is reserved for spontaneous neural activity, at least before birth and in early development (Katz & Shatz, 1996). Spontaneous activity as it occurs in the developing retina, cochlea, spinal cord, cerebellum and hippocampus, among others, is patterned. It provides important signals for the development of neurons and their connections (Blankenship & Felller, 2010). A key example is retinal wave activity (Galli & Maffei, 1988; Luhmann, 2022; Shatz, 1996). It arises in the interaction of amacrine and retinal ganglion cells (Xu et al., 2016) and occurs spontaneously, including a period before any visual input is available (Maccione et al., 2014). This is because the ganglion cells precede the other retinal cells, in particular the photoreceptors, in development (D’Souza, & Lang, 2020).

Retinal waves play a decisive role in the formation of the topographic map structure of the retina (Arroyo & Feller, 2016), As the sole retinal output units, retinal ganglion cells project the retinal waves onto V1 (Alexander et al 2011), and to the visual system beyond (Ackman et al., 2012; Xu et al., 2016). The neurophysiological mechanisms of retinal wave propagation effects are complex; involving active signaling and diffusion within the neural architecture (Leybaert & Sanderson, 2012). Retinal waves occur in three stages of neural development, evoked by different forms of neurotransmission (Maccione et al., 2014; Xu et al., 2016).

A basic question, however, has remained underexplored: how the retinal ganglion cells could develop into retinal output hubs in the first place. This question is addressed here within the framework of adaptive rewiring models (Gong & van Leeuwen, 2003; 2004; Jarman et al., 2017; Rentzeperis et al., 2022; Li et al., 2023). Using highly simplified neural network models, adaptive rewiring aims to understand in a principled manner the emergence of complex large-scale structures of the brain, i.e. modular small-world networks (Bullmore & Sporns, 2009) with rich club effect (Zamora-López, et al., 2010; van den Heuvel & Sporns, 2011). In adaptively rewiring networks with ongoing oscillatory (Gong & van Leeuwen, 2003; 2004) or spiking (Kwok et al, 2007) activity, the connectivity structure is updated continuously by establishing new connections between units that interact strongly, while underused ones are cut. This optimizes the communication efficiency of the networks, leading to modular small-world structures (van den Berg & van Leeuwen, 2004; Rubinov et al., 2009) with a rich-club effect (Hellrigel et al., 2019).

Subsequent adaptive rewiring studies have simplified modeling even further, by representing the flow of activity through random walks on a graph. This enabled closed-form specification of the activity flow in terms of network diffusion kernels (Jarman et al., 2017; Rentzeperis et al., 2021). A recent move towards greater neural realism takes into account that neural connections are preferentially formed locally (Antonello et al., 2022). This spatial preference was initially modeled by postulating wiring costs (Jarman et al., 2014). A large proportion of connections in the brain is formed by calcium signals percolating through gap junctions between spatially neighbouring neurons (Hettiarachchi et al., 2010, Weiss et al., 2022). Therefore, later models introduced a special rewiring rule based on spatial proximity alone (Calvo Tapia et al., 2020; Li et al., 2023).

Following earlier work by Rentzeperis et al. (2022), Li et al. (2023) showed that adaptive and spatial rewiring jointly enable the formation of convergent-divergent units (Shaw et al., 1982; Kumar et al., 2010). Convergent-divergent units are network structures comprising one or more convergent hubs, which collect input from a range of local neurons, and feed their signals to a relatively isolated set of units (a module), which in turn feeds a divergent hub, which broadcasts the output regionally or globally (Figure 1A). These units support neural signal flow and enable context-sensitive neural computation. Key examples of such units are the context-sensitive circuitry of V1: activity converges on orientation-selective neurons in layers 2/3 that send their input to somatostatin (SOM) cells. These act as divergent hubs to broadcast their response back to the network (Adesnik et al., 2012; Niell & Scanziani, 2021), thereby enhancing orientation selectivity of V1 neurons (Song et al., 2020).

**Figure 1.**
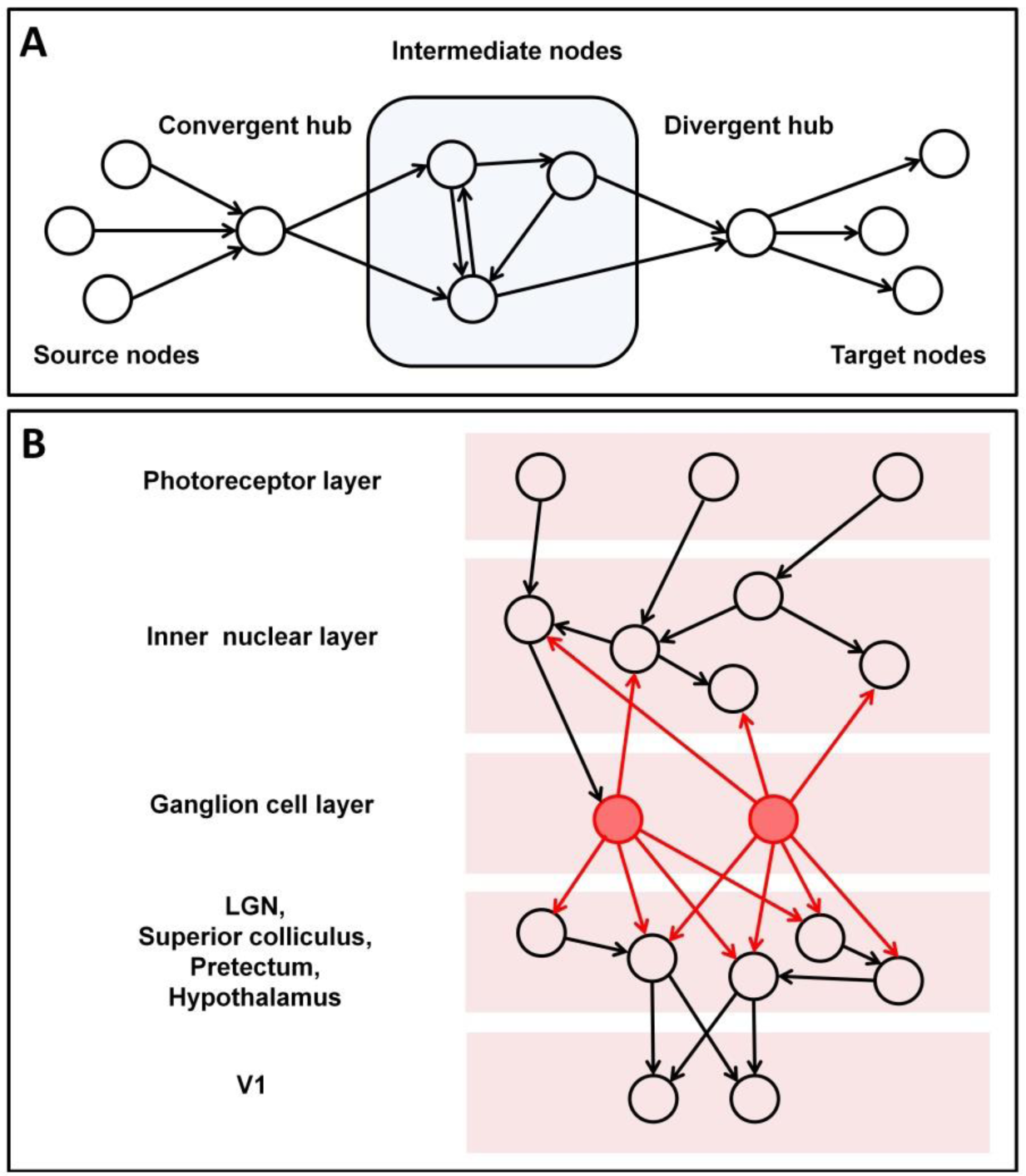
Schematic network representations. **A.** From Li et al., 2023. Schema of a convergent-divergent unit. In a convergent-divergent unit, a convergent hub collects inputs and passes the information to a divergent hub through a subnetwork of intermediate nodes. The nodes sending information to the convergent hub are referred as source nodes, and those receiving information from the divergent hub as target nodes. Note that typically the source and target nodes can show overlap, i.e., a node can be both a source and a target node. **B.** Schema of retinal circuitry. Photoreceptors feed their input into the rest of retinal cell types. Out of these, retinal ganglion cells (RGCs) act as divergent hubs that propagate their activity throughout the network, serving as input to other parts of the brain, such as the superior colliculus, the pretectum, the hypothalamus and, importantly, the Lateral Geniculate Nuclei (LGN), that connect with V1.

Divergent hubs also exist in the retina. Retinal ganglion cells (RGC) engage in reciprocal connections with amacrine cells in the inner nuclear layer and other retinal cells and are responsible in particular for the propagation of visual stimuli from both eyes to V1 as well as a number of other brain structures. Their axons leave the retina via the optic disk, joining to form the optic nerve en route to the optic chiasm, where about 60 % of them cross over to the other side, while the rest remains uncrossed. From here, the fiber bundles, which now each contain RGC axons from both eyes, continue as the optic tract, to terminate in the superior colliculus, the pretectum, the hypothalamus, and the Lateral Geniculate Nuclei (LGN) of the thalamus (Purves et al., 2001). The superior colliculus and the LGN, from which the visual signals ultimately reach V1 (Figure 1B), may be regarded as convergent hubs in downstream convergent-divergent units. In other words, even before V1 is reached, RGCs broadcast the visual signal to a number of brain regions. RGCs are responsible for distinct key visual computations, such as direction selectivity during motion processing, contrast detection and orientation selectivity among others (see Kerschensteiner, 2022 for a review).

### Ganglion cells as divergent hubs

The role of RGC as divergent hubs is the main focus of our current study. The self-organizing networks of Li et al. (2023) lack facilities for receiving sensory input. Previous studies (Haqiqatkhah & van Leeuwen, 2022) showed that input systems could develop, and reach a degree of segregation in their entirety from the central network structure, without however destroying the integrity of the network. These studies did not offer suggestions on how retinal wave activity could constrain the evolving architecture of the visual system, though. We consider, in particular, whether retinal wave activity could play a decisive role in developing retinal ganglion cells into divergent hubs. We also consider whether this is achieved without hampering the developing complexity of the rest of the network, in particular, the emergence of convergent-divergent units.

### Model strategy

For our model, we adopt the activity-based adaptive rewiring and spatial-proximity based rewiring rules as proposed by Li et al. (2023). A third rule also used in Li et al. (2023), alignment, was not included here because its effect on network structure was found to be negligible. Adaptive rewiring in Li et al. (2023) is based on signal flow represented as network diffusion. Incoming diffusion for each node was determined according to a consensus criterion (Ren et al., 2007) and outgoing diffusion was determined according to advection (Chapman, 2015). A proportion of the times, adaptive rewiring was based on incoming flow, and a proportion on outgoing flow. In addition, a proportion of the times spatial rewiring was used, replacing a spatially distant connection with a spatially local one.

Initially random networks develop into complex networks according to iterative application of these rules. A subset of the nodes is designated to represent retinal ganglion cells. Retinal waves in ganglion cells are initiated in interaction with amacrine cells (Xu et al., 2016). We modeled this effect by supplying nodes designated as ganglion cells for a given proportion of times throughout the rewiring process with increased outgoing flow. We observe whether these nodes develop into divergent hubs. In addition, we consider the evolution of convergent-divergent units, and the participation of these designated nodes therein.

We find that over successive rewirings of an initially random network, as in Li et al. (2023), a complex network structure evolved that was spatially local, small-world, modular, and contained convergent-divergent units with both convergent and divergent hub nodes. A significant proportion of the nodes designated as ganglion cells developed into divergent hubs. The remaining hubs in the network developed spontaneously. This has the relevant implication that the formation of ganglia as divergent hubs in evolving complex networks can be stochastically controlled by the retinal waves themselves.

## Method

### Notation and definitions

In accordance with Li et al. (2023), we define a directed graph (digraph) as the set *G* = (*V*, *E*, *W*), where *V* = {1,2,…,*n*} represents the set of nodes, *E* ⊂ *V* × *V* denotes the set of ordered pairs of nodes, in which (*j*, *i*) ∈ *E* indicates directed edges from node *j* to *i* (*j* → *i*), and *W* = {*w*_*ij*_: *i*, *j* ∈ *V*} represents the nodes’ weights. The adjacency matrix, *A* = [*A*_*ij*_]_*i*, *j* ∈ *V*_, with dimensions *n* × *n*, contains these weights, where *A*_*ij*_ = *w*_*ij*_, with *w_*ij*_* > 0 when (*j*, *i*) ∈ *E* while the length of *j* → *i* is given by 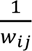. When (*j*, *i*) ∉ *E*, *w*_*ij*_ = 0. |*V*| = *n* and |*E*| = *m*, respectively, denote the numbers of nodes and directed edges. Note that the weights are drawn from a probability distribution and will remain fixed, except when the retinal wave rule applies.

We define the in-links of a node **i** ∈ **V** as the set of edges directed towards *i*, and the out-links as the set of edges originating from *i*. The in-degree neighbourhood of *i*, *N*_*in*_(*i*), comprises the tails of the in-links connected to *i*, while the remaining nodes, *V* − *N*_*in*_(*v*), are denoted as 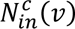. The in-degree of a node i is determined by the count of its in-links. Similarly, the out-degree neighbourhood of *i*, denoted as *N*_*out*_(*i*), consists of the heads of the out-links emanating from *i*, while 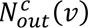 represents the remaining nodes. The out-degree of a node *i* is calculated as the number of its out-links. Considering an ordered pair of nodes (*u*, *v*), a directed walk from *u* to *v* is a sequential list of edges {(*i*_0_, *i*_1_), (*i*_1_, *i*_2_),…,(*i*_*k*−1_, *i*_*k*_): *i*_0_ = *u*, *i*_*k*_ = *v*, (*i*_*k*−1_, *i*_*k*_) ∈ *E*} (Bender & Williamson, 2010). A directed walk is described as a directed path if all the vertices along it are distinct.

### Consensus and advection dynamics

The consensus and advection kernels represent the communication intensity between nodes, and are expressed as:

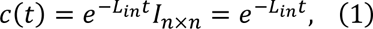

and:

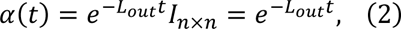

where *c*(*t*) and *a*(*t*) are the consensus and advection kernels respectively. (See Appendix A, “Derivation of the consensus and advection kernels”).

### Rewiring Rules

Our investigation focuses on analyzing the network’s structural changes through iterative rewiring of its edges. During each iteration, a node *v* ∈ *V* is randomly chosen for rewiring. If we are to rewire one of the in-links from *v*, an existing edge (*k*, *v*) ∈ *E* will be removed, and a new edge (*l*, *v*) ∉ *E* will be added, connecting *v* to a node *l* that was not previously linked to *v*. Similarly, when rewiring an out-link of *v*, we replace an existing edge (*v*, *k*) ∈ *E* with a new edge (*v*, *l*) ∉ *E*. To determine the selection of nodes *k* and *l* during each rewiring step, we employ one of the following principles, each with a fixed probability: the adaptive rewiring principle or the proximity-based principle.

#### Adaptive rewiring

Adaptive rewiring implies that an underutilized connection is eliminated while a new connection is established between two previously unconnected nodes that exhibit the most intense flow between them, considering all indirect paths. A variety of network topologies emerge when rewiring the in-degree neighbourhood using the consensus algorithm or rewiring the out-degree neighbourhood using the advection algorithm (Rentzeperis et al., 2022). When rewiring in-links we employ the consensus kernel, and rewiring of out-links uses the advection kernel to estimate the communication intensity for rewiring the nodes.

When rewiring an in-link of node *v*, *k* is the node in *N*_*in*_(*v*) where (*k*, *v*) has the lowest value in the consensus kernel, denoted as *k* = *argmin*_*u*∈*Nin*(*v*)_{*c*(*t*)_*vu*_}. Similarly, *l* is the node in 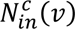 such that (*v*, *l*) has the highest value in the consensus kernel, expressed as 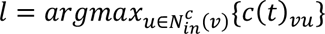. When an out-link is rewired, *k* is the node in *N*_*out*_(*v*) such that (*v*, *k*) has the lowest value in the advection kernel, indicated by *k* =*argmin*_*u*∈*Nout*(*v*)_{*a*(*t*)_*uv*_}. Correspondingly, *l* is the node in 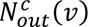 that makes (*l*, *v*) have the highest value in the advection kernel, and is represented as 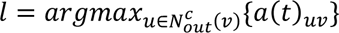. Whether rewiring an in-link or an out-link of node *v*, *l* is the node chosen for rewiring in both cases.

Both the consensus and advection kernels incorporate a time variable, *t*, set to a predetermined parameter known as the rewiring rate τ. This parameter signifies the duration between successive rewiring iterations (τ adopted a fixed value of 1 during our simulations).

#### Retinal wave rule

This rule is a special case of adaptive rewiring. The otherwise constant out-link weights of a node *c*, to which we refer as *wave initiator* node, are boosted by adding a constant value, *B*, to their weights. A fixed value of *B* = 1 was adopted for our simulations. The boost represents the event that node *c* is releasing connection-forming Ca^2+^ in response to retinal wave activity. The weights from the entire network are then renormalized by dividing their values by the sum of all the weights. Next, rewiring takes place according to the adaptive rewiring rule. Afterward, all the weights, including those of node *c*, are reset to their original values, prior to adding the boost.

#### Proximity-based rewiring

For this, we assume the digraph to be embedded within a two-dimensional Euclidean space. Each node *i* is denoted as 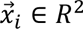. In this space, proximity-based rewiring means that the longest connection is removed and substituted with the spatially shortest link between two nodes that were previously unconnected. The distance between node *i* and node *j* is calculated using 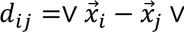, where ∨·∨ represents the Euclidean distance. When rewiring an in-link of node *v*, *k* is the node in *N*_*in*_(*v*) that has the greatest distance from *v*; that is, *k* = *argmax*_*u*∈*Nin*(*v*)_{*d*_*uv*_}. Similarly, *l* is the node in 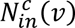 that has the shortest distance from 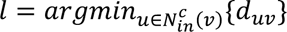. Conversely, when rewiring an out-link of node *v*, *k* is *argmax*_*u*∈*Nout*(*v*)_{*d*_*vu*_}, and *l* is 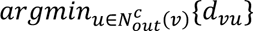.

### Rewiring algorithm

For simplicity, the rewiring process maintains a constant number of nodes and edges within the networks. Other possibilities such as growing networks (Gong & van Leeuwen, 2003) and pruning networks (van den Berg et al., 2012) have also been investigated. To initiate the rewiring process, a random directed network *D* = (*V*, *E*, *W*) is established, with a given number of nodes, *n*, and edges, *m*. The *m* edges are assigned to node pairs, selected randomly without replacement from all possible *n*(*n* − 1) node pairs. Weights drawn from a normal probability distribution, *N*(1, 0.25^2^), are assigned at random to the edges. In the extremely unlikely event that a negative weight is sampled (3.17 × 10^−5^ probability), it is set to 0.05.

The iterative rewiring process follows as outlined below:

- *Step 1:* Randomly choose a node *v* ∈ *V* such that its in-degree is not zero and not equal to *n* − 1. Alternatively, choose a random node *v* ∈ *V* with an out-degree that is not zero and not equal to *n* − 1.
- *Step 2:* Determine whether to rewire an in-link or an out-link of *v* based on a probability value *p*_*in*_. If *p*_*in*_ is set to 1, only in-links are subject to rewiring, resulting in an iterative process known as in-link rewiring. Conversely, if *p*_*in*_ is set to 0, the iterative process is referred to as out-link rewiring (*p*_*in*_ adopted a fixed value of 0.5 during our simulations).
- *Step 3:* Depending on the outcome of Step 2, select a random in-link or out-link of *v* and rewire it using one of the following rewiring rules: adaptive, retinal wave (a special case of adaptive rewiring), or proximity rewiring. The probabilities of selecting each rule are denoted as *p*_*adaptive*_, *p*_*wave*_, and *p*_*proximity*_, respectively.
- *Step 4:* Repeat Steps 1 to 3 until a total of *M* edges have been rewired.

Network evolution was tested for different probabilities of *p*_*adaptive*_, *p*_*wave*_, and *p*_*proximity*_. For each set probability, 150 initially random networks were evolved throughout 4000 iterations. Their resulting structure was analysed using measures of graph topology and the results were averaged across the 150 networks. Parameter *p*_*in*_ (i.e. the probability that either an in-link or an out-link is rewired) was set to a value of 0.5, τ (i.e. the duration between successive rewiring iterations) was fixed to 1 and *B* (i.e.: strengthening of the out-link weights of a wave initiator node during wave occurrence) had a value of 1.

The following measures were used:

### Modularity

Modularity gauges the degree to which a network can be divided into distinct communities. In the context of a weighted digraph, we adopt the modularity definition proposed by Arenas et al. (2007):

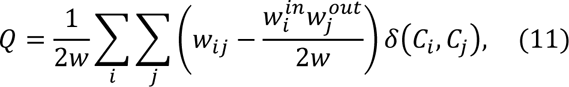

where 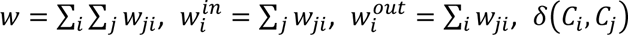 designates the Kronecker delta function and *C*_*i*_ denotes to which community node *i* belongs to. We use the algorithm by Leicht & Newman (2008) to find these communities.

### Average efficiency

The average efficiency metric measures how effectively information is transmitted across a network. It is calculated as the average of the reciprocals of the shortest directed path lengths between all pairs of nodes (Latora & Marchiori, 2001). Despite the networks being spatially embedded, we focuss on assessing network efficiency concerning its connectivity structure and weights. We use the topological distance, 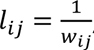, which serves as an indicator of transmission complexity (Opsahl et al., 2010). At the neuronal level, a stronger synapse (large *w*_*ij*_) facilitates the easier transmission of electric nerve impulses between two neurons, resulting in smaller values of *l*_*ij*_.

For a given pair of nodes (*u*, *v*), a directed walk from *u* to *v* consists of an ordered sequence of edges {(*i*_0_, *i*_1_), (*i*_1_, *i*_2_),…, (*i*_*k*−1_, *i*_*k*_): *i*_0_ = *u*, *i*_*k*_ = *v*, (*i*_*k*−1_, *i*_*k*_)ϵ*E*} (Bender & Williamson, 2010). A directed walk is a directed path when all its vertices are different. Average efficiency is defined as:

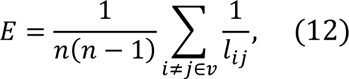

where *l*_*ij*_ represents the length of the shortest directed path from node *i* to node *j*, signifying the most straightforward transmission route between the two nodes. In those cases where there is no transmission route from *i* to *j*, *l*_*ij*_ = ∞.

### Number of connected node pairs

A connectedness measure for a directed graph (digraph) is based on ordered pairs, (*i*, *j*), which are considered connected if there exists a directed path from node *i* to node *j*. The count of such connected node pairs serves as an indicator of the level of information exchange within the digraph. The upper limit for the number of connected node pairs is *n*^2^, and occurs when every node can transmit information to any other node, including itself. We use this metric to quantify the degree of connectedness within a given digraph.

### Convergent and divergent hubs, convergent-divergent units

We classify nodes as convergent hubs if they have at least one out-link and a number of in-links that surpasses a given threshold. These hubs serve as a substrate to collect dispersed information. Conversely, divergent hubs are nodes that possess at least one in-link and a number of out-links that exceed a predefined threshold. These hubs serve to disseminate information widely. We set the threshold for both convergent and divergent hubs to the value of 15 in all evaluations. We consider a convergent-divergent unit to be formed if a convergent and divergent hub are linked by one or more directed paths.

## Results

### Modularity, average efficiency and connectedness are preserved in networks with retinal waves

We assessed the average degree of modularity, efficiency and connectedness in the resulting networks after 4,000 iterations. Thishe number roughly represents the period of the first three postnatal weeks, in which synapses develop in the mouse retina (Hoon et al., 2014). The modularity indices in Figure 2 (left) show the extent to which the networks have a modular structure. Modularity increased with the proportion of proximity-based rewiring. Average efficiency and connectedness (Figure 2, middle and right, respectively) showed similar trends, suggesting that network efficiency is mostly related to its degree of connectedness. The occurrence of retinal waves did not interfere with the emergence of modular, efficient and connected networks.

**Figure 2.**
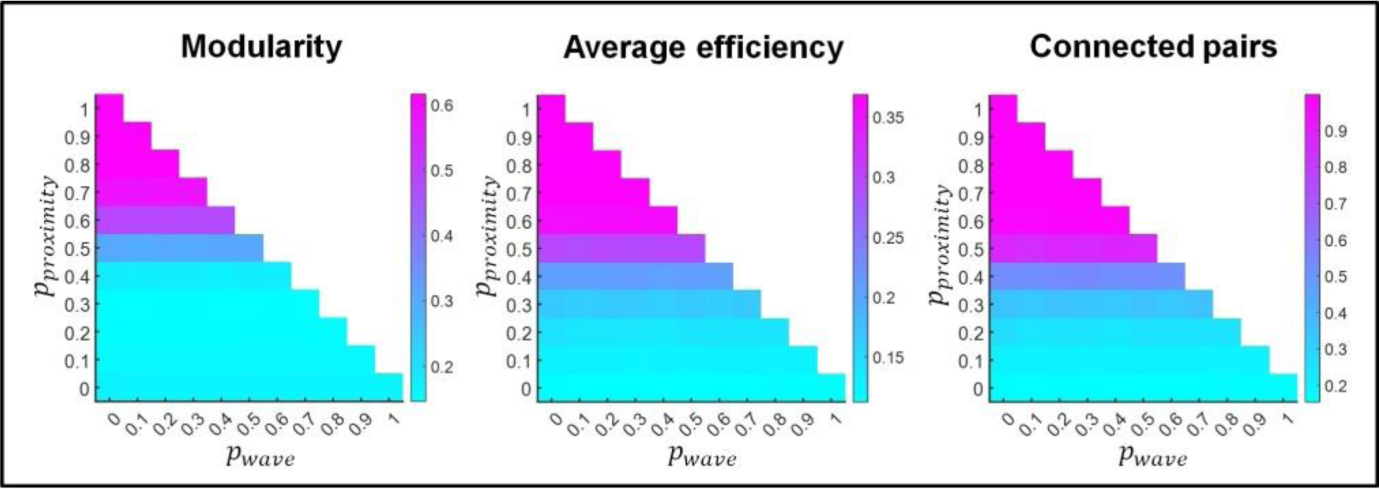
Modularity index (left), average efficiency (middle) and proportion of connected node pairs (right) of the networks resulting from rewiring. Results are shown as a function of the probability of proximity-based rewiring and retinal wave-based rewiring. The remaining probability (i.e. when the sum of the probabilities for proximity-based and retinal wave-based rewiring does not reach 1) corresponds to adaptive rewiring based on graph diffusion. For each combination of rewiring principle probabilities (e.g.: *p*_*proximity*_ = 0.5, *p*_*w*__*ave*_ = 0.2, *p*_*adaptive*_ = 0.3), results are obtained by averaging over 150 networks that evolve from 150 initially random networks.

### Increased number of out-links in retinal wave initiator nodes

We investigated to what extent retinal wave initiator nodes developed into divergent hubs. Their number of out-links was compared to that of the other, non-initiator nodes. By computing the ratio of their out-links, we obtained a metric of the consistency of wave initiator out-link development. The consistency was found to increase with an increasing proportion of retinal wave-based rewiring. For the non-initiator nodes, a minor tendency in the opposite direction was observed (i.e. the more seldom the waves, the larger the number of out-connections in non-initiator nodes; see Figure 3A). Indices analogous to those in Figure 3A were calculated for in-links (Figure 3B). Wave initiator nodes developed a lower number of in-links with larger proportions of retinal wave-based rewiring, whereas for non-initiator nodes in-link connectivity showed there was a minor tendency in the opposite direction.

**Figure 3.**
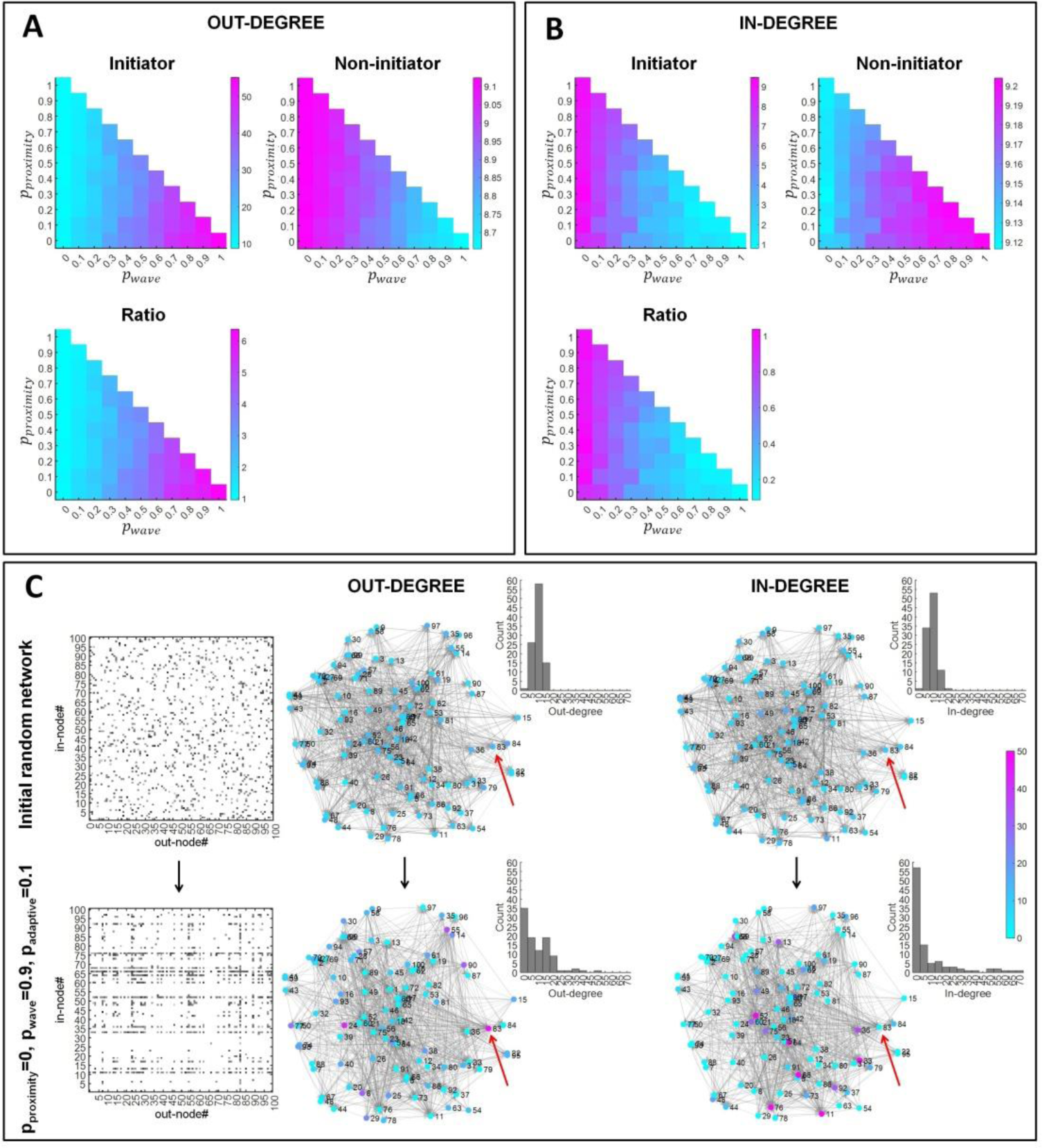
Out-link and in-link connectivity of retinal wave initiator and non-initiator nodes. **A & B.** In each cell (cell A & cell B), the left upper panels represent the average number of out-connections (A) and in-connections (B) in retinal wave initiator nodes. The right upper panels represent the same for non-initiator nodes. Note that the vertical scales differ between left and right panels within the top cells. The lower panels represent the ratio between the two upper panels, indicating the probability regions for the different rewiring principles where retinal wave initiator nodes most consistently develop a significant number of out-connections (A) and in-connections (B). Results are shown as a function of the probability of proximity-based rewiring and retinal wave-based rewiring. The remaining probability (i.e. when the sum of the probabilities for proximity-based and retinal wave-based rewiring does not reach 1) corresponds to adaptive rewiring based on graph diffusion. For each combination of rewiring rule probabilities (e.g.: *p*_*proximity*_ = 0.5, *p*_*wave*_ = 0.2, *p*_*adaptive*_ = 0.3), results are obtained by averaging over 150 networks that evolve from 150 initially random networks. **C.** Example network structure of an initially random network (upper plots) and a network after rewiring using *p*_*proximity*_ = 0, *p*_*wave*_ = 0.9, *p*_*adaptive*_ = 0.1 (lower plots). The leftmost plots show each of the adjacency matrices, indicating the out-connectivity and in-connectivity of each node, according to its identifier number (node#). The plots on their right show the spatially embedded networks. The numbers next to the nodes indicate the node identifier numbers, and the number of out-connections (OUT-DEGREE) and in-connections (IN-DEGREE) of each is expressed according to the colour bar. The retinal wave initiator is node number 83 (indicated by a red arrow). For each network, the histograms depict the distributions of, respectively, the nodes’ in and out-degree.

Figure 3C (left) illustrates how an initially random network evolves with a high probability of wave-based rewiring (*p*_*wave*_ = 0.9). The wave initiator node (Node 83) develops the largest number of out-links amongst the nodes. Figure 3C (right) shows the in-link connectivity for the same network as in Figure 3C (left). The wave initiator node does not develop a large number of in-links. The in- and out-degree histograms in Figure 3C evidence the changes in network connectivity structure as a result of rewiring. See Figure S1 in the Supplementary material, “Network embeddings”, for more examples of network topologies resulting from different probabilities for **p*_*proximity*_, *p*_*wave*_*and **p*_*adaptive*_*.

An important question is whether retinal waves interfere with the formation of convergent-divergent units. We first show the proportion of convergent and divergent hubs that exist in the networks resulting from rewiring (Figure 4A and 4B respectively). Both convergent and divergent hubs most likely formed with intermediate proximity-based rewiring probabilities (i.e. in the range 0.4-0.5). In Figure 3A we show that for the largest wave-based rewiring probabilities, initiator nodes consistently developed the largest number of out-links. However, in these cases, initiator nodes also received the smallest number of in-links (see Figure 3B). To qualify as a divergent hub, a node needs to have an elevated number of out-links, but also at least one in-link. This criterion was rarely met when wave-based rewiring had the largest probability, and thus, retinal wave initiator nodes did not often become divergent hubs in this case. For this reason, the greatest likelihood of finding initiator nodes developing into divergent hubs was obtained in the intermediate regime. Figure 4C (upper panel) shows that initiator nodes very likely became divergent hubs (i.e. probabilities around 0.6) when proximity-based rewiring was in the probability range 0.5-0.8 and wave-based rewiring was in the range 0.2-0.5.

**Figure 4.**
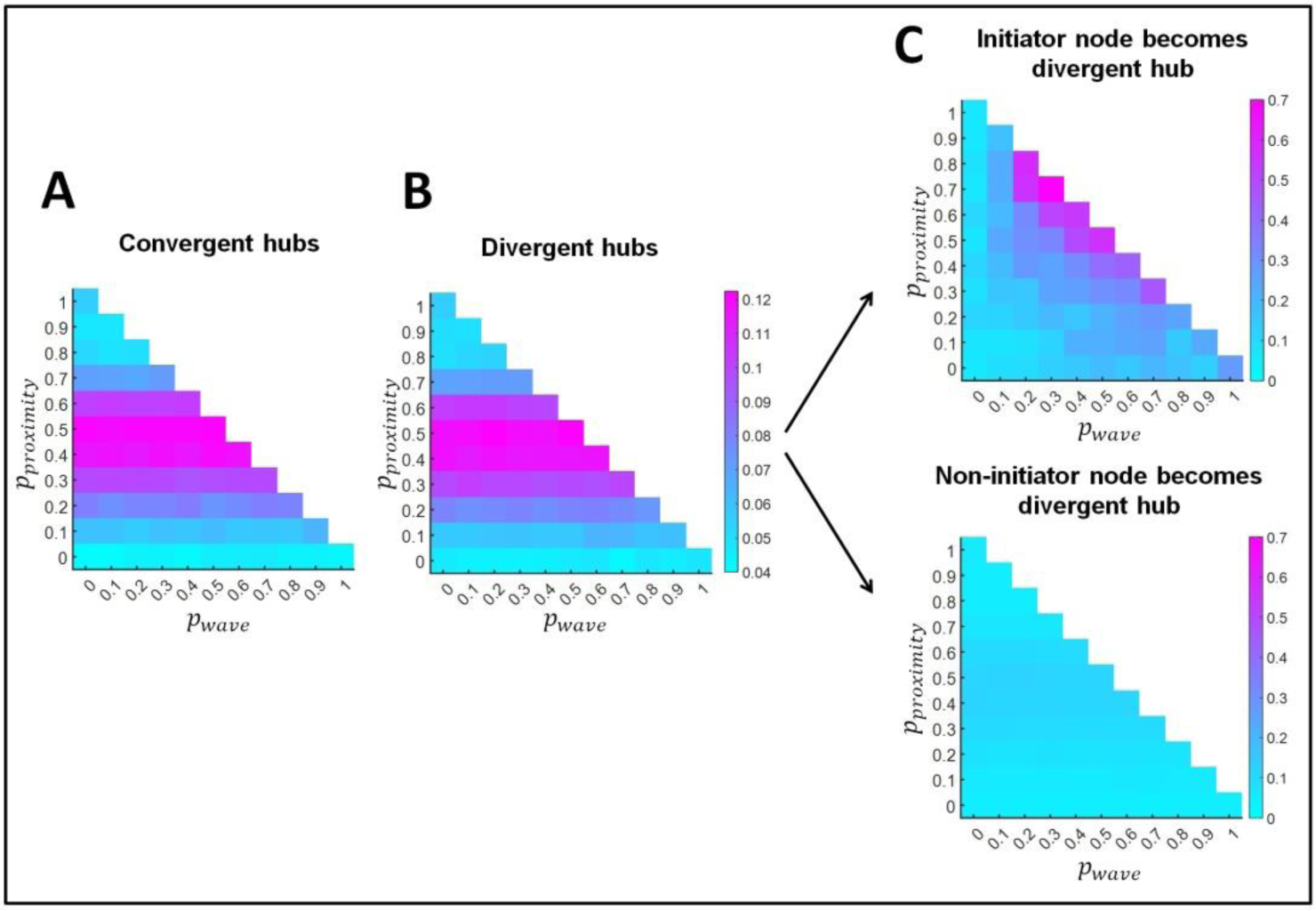
Formation of convergent and divergent hubs. **A.** Proportion of convergent hubs. **B.** Proportion of divergent hubs. **C. Upper panel**. Proportion of network instances where retinal wave initiator nodes become a divergent hub. **Lower panel**. Out of 1000 random selections of nodes that are not retinal wave initiators, proportion of times that these are a divergent hub. Results are shown as a function of the probability of proximity-based rewiring and retinal wave-based rewiring. The remaining probability (i.e. when the sum of the probabilities for proximity-based and retinal wave-based rewiring does not reach 1) corresponds to adaptive rewiring based on graph diffusion. For each combination of rewiring rule probabilities (e.g.: *p*_*proximity*_ = 0.5, *p*_*w*__*ave*_ = 0.2, *p*_*adaptive*_ = 0.3), results are obtained by averaging over 150 networks that evolve from 150 initially random networks.

On the other hand, when we randomly selected 1000 non-initiator nodes and tested whether they were divergent hubs, we evidenced a much lower probability than that of initiator nodes for becoming divergent hubs (Figure 4C, lower panel). This probability was maximal when proximity-based rewiring was in the range 0.4-0.5.

Now that we have demonstrated the formation of convergent and divergent hubs, we proceed to the formation of convergent-divergent units, which result from the existence of convergent and divergent hubs linked through a direct path. Figure 5A shows the average total number of convergent-divergent units that evolved in the networks. Not surprisingly, its trend is similar to that of convergent and divergent hubs. Figure 5B shows the success rate for the formation of convergent-divergent units. We define this success rate as the proportion of network instances (out of the 150 networks resulting from rewiring) where at least one convergent-divergent unit formed. The success rate increased with larger probabilities for proximity-based rewiring, as also did the connectedness of networks (Figure 2, rightmost panel). That is, the success rate of forming convergent-divergent hubs depends, not only on the number of convergent and divergent hubs, but also on the number of direct paths between nodes.

**Figure 5.**
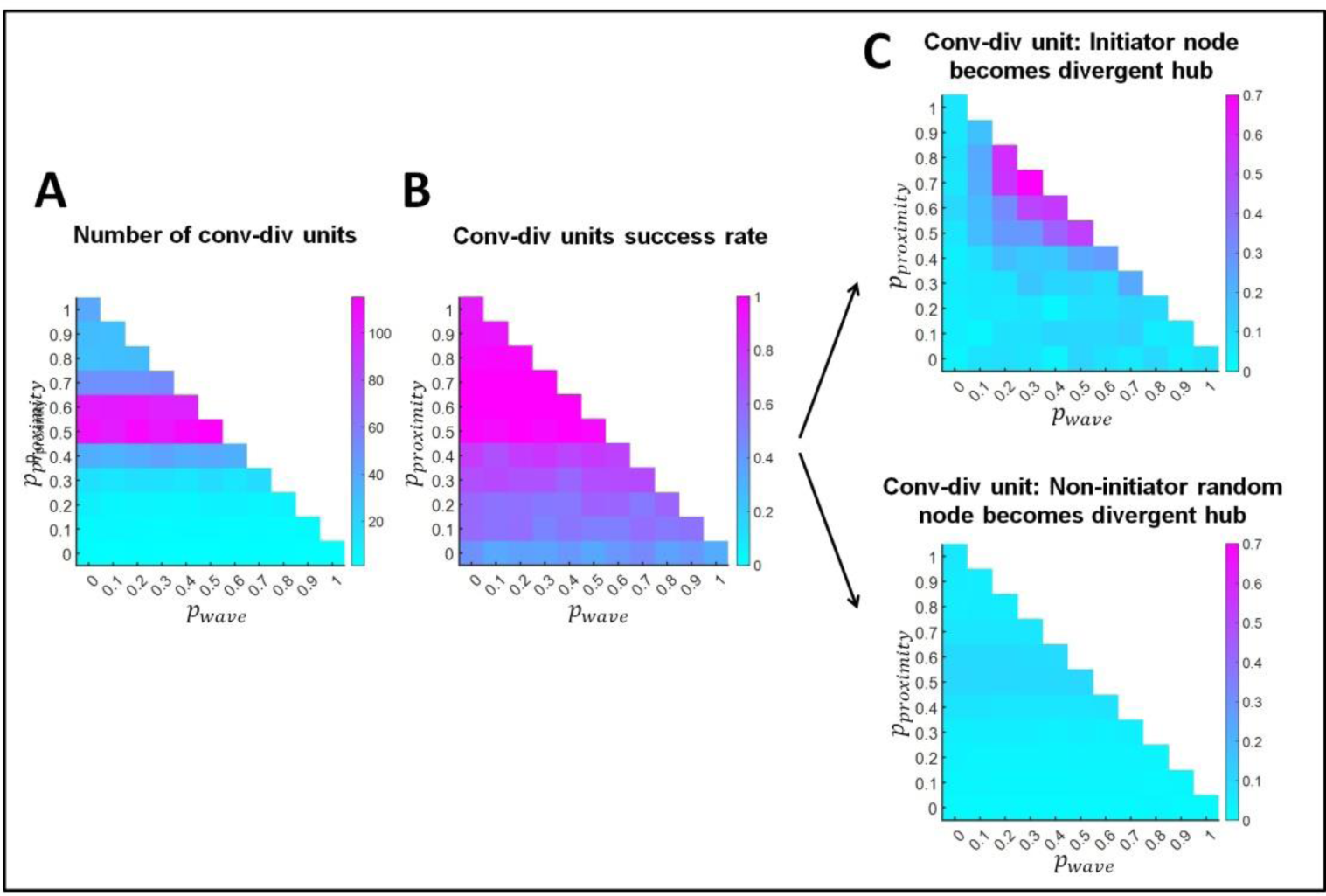
Formation of convergent-divergent units. **A.** Total number of convergent-divergent units. **B.** Proportion of network instances where at least one convergent-divergent unit exists. **C. Upper panel**. Proportion of network instances where a retinal wave initiator node becomes the divergent hub of a convergent-divergent unit. **C. Lower panel.** Out of 1000 random selections of nodes that are not retinal wave initiators, proportion of times that these are a divergent hub of a convergent-divergent unit. Results are shown as a function of the probability of proximity-based rewiring and retinal wave-based rewiring. The remaining probability (i.e. when the sum of the probabilities for proximity-based and retinal wave-based rewiring does not reach 1) corresponds to adaptive rewiring based on graph diffusion. For each combination of rewiring rule probabilities (e.g.: *p*_*proximity*_ = 0.5, *p*_*wave*_= 0.2, *p*_*adaptive*_ = 0.3), results are obtained by averaging over 150 networks that evolve from 150 initially random networks.

We next considered whether wave initiator nodes were more likely than non-initiator nodes to become the divergent hub of a convergent-divergent unit. Figure 5C (upper panel) indicates that initiator nodes more likely became divergent hubs given a specific combination of probabilities for the rewiring principles (i.e. 0.2-0.5 probability for retinal wave-based rewiring and 0.5-0.8 probability for proximity-based rewiring). Within this range, the probability that a retinal wave initiator node would become a divergent hub increased with the retinal wave-based rewiring probability. The probability that a non-initiator node would become a divergent hub in the convergent-divergent unit was calculated across 1000 random selections of non-initiator nodes. Importantly, the probability of becoming the divergent hub was larger for retinal wave initiator nodes than for non-initiator nodes in the convergent-divergent unit (see Figure 5C, upper and lower panels).

In convergent-divergent units, nodes on directed paths between the divergent and the convergent hub are referred to as intermediate nodes. The subgraph integrated by these nodes is called the intermediate subgraph, and processes the information from the convergent hub. The extent to which this subgraph is isolated from the rest of the network determines its processing style and context sensitivity. For intermediate subgraphs containing more than one node, Figure 6 shows their size (i.e. number of nodes, *n*) and density (number of edges, *m*, divided by the maximum number of edges possible: *m*⁄[*n* (*n* − 1)⁄2]). We calculated these values both for networks where a retinal wave initiator node was the divergent hub (Figure 6A) and networks where the divergent hub was a non-initiator node (Figure 6B). In both cases, the size of the intermediate subgraph increased with the probability of proximity-based rewiring. Regarding density, it decreased with increasing proximity-based rewiring probability, until *p*_*proximity*_ > 0.5, where a floor value was reached. This can easily be explained by the fact that in the reported probability regions, all nodes of the graph, excluding the convergent and divergent hubs, were part of the intermediate subgraph.

**Figure 6.**
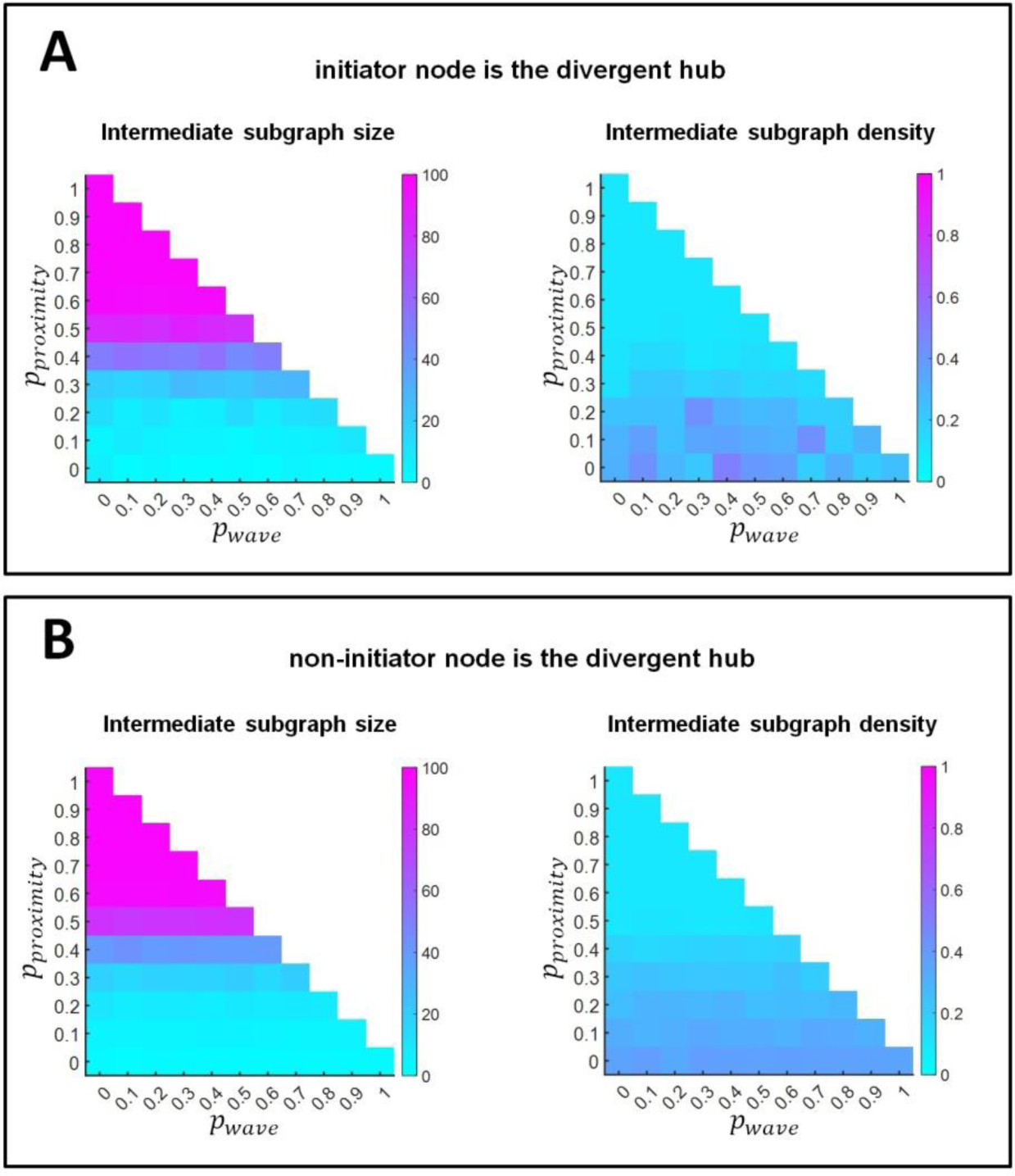
Subgraph structure in convergent-divergent units. **A.** Convergent-divergent units where retinal wave initiator nodes are the divergent hubs. **B.** Convergent-divergent units where non-retinal wave initiator nodes are the divergent hubs. The left panels represent the size, as number of nodes, of the intermediate subgraphs and the right panels represent their density. Results are shown as a function of the probability of proximity-based rewiring and retinal wave-based rewiring. The remaining probability (i.e. when the sum of the probabilities for proximity-based and retinal wave-based rewiring does not reach 1) corresponds to adaptive rewiring based on graph diffusion. For each combination of rewiring rule probabilities (e.g.: *p*_*proximity*_ = 0.5, *p*_*w*__*ave*_ = 0.2, *p*_*adaptive*_ = 0.3), results are obtained by averaging over 150 networks that evolve from 150 initially random networks.

### Increased number of in-links in nodes targeted by retinal wave initiator nodes

Interestingly, in the lower panel in Figure 3B we observed that while non-initiator cells lose outgoing connectivity, they gain incoming connectivity. We might consider this as an effect of renormalization of the network weights. In that case, the effect would be equally distributed among all non-targeted nodes. However, it rather affects in particular the nodes targeted by the out-links of initiator nodes. Figure 7B shows the in-degree of nodes targeted by the out-connections of retinal wave initiator nodes as well that of non-targeted nodes. The ratio is displayed in the lower panel of Figure 7B.

**Figure 7.**
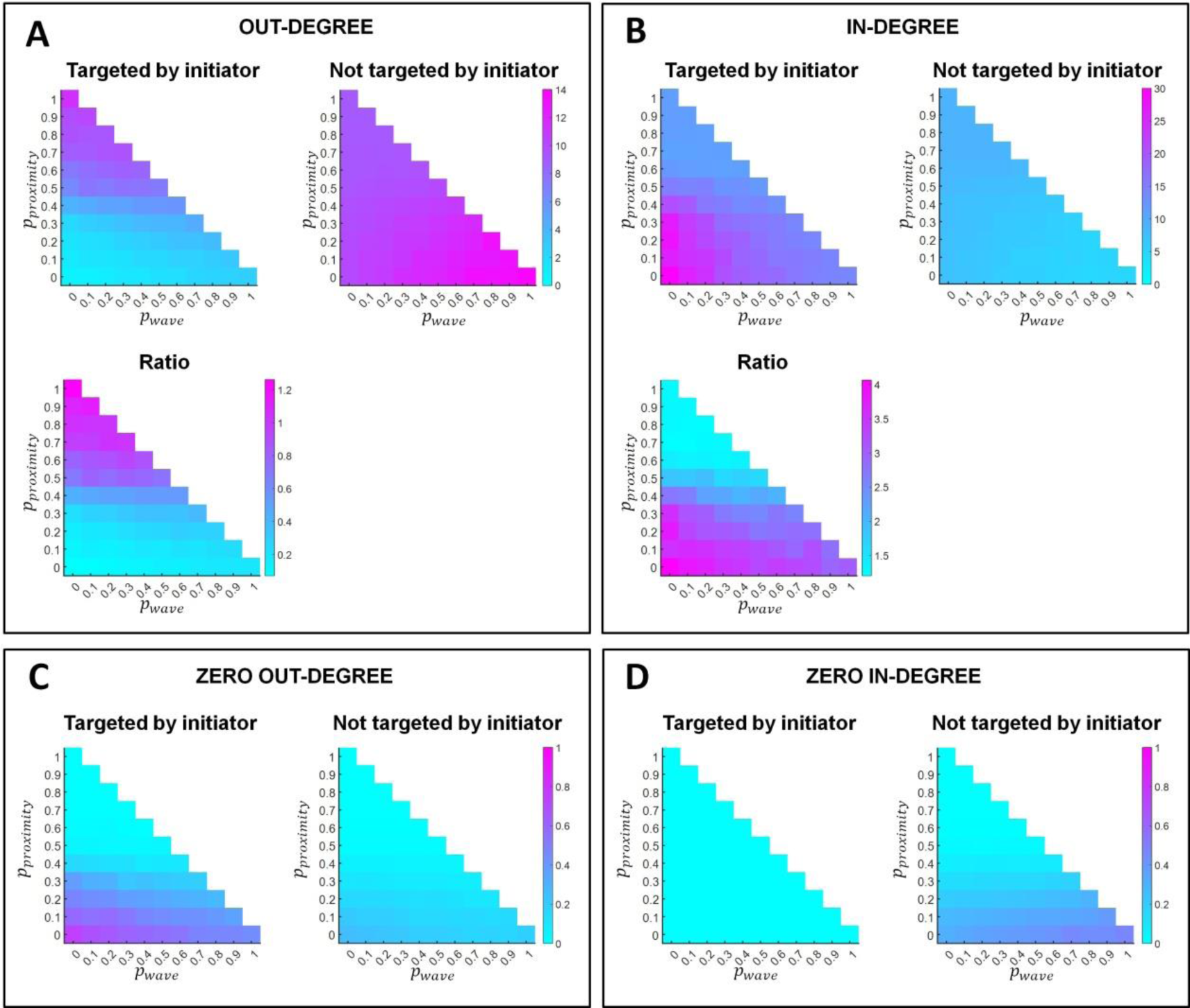
In and out connectivity of nodes targeted by the outlinks of retinal wave initiator nodes and the rest of non-targeted nodes. **A & B.** In each cell (cell A & cell B), the left upper panel represents the average number of out-connections (A) and in-connections (B) of nodes targeted by initiators (left panel) and not targeted by initiators (right panel). Note that the vertical scales differ between left and right panels within the top cells. The lower panels represent the ratio between the two upper panels, indicating the probability regions for the different rewiring principles where nodes targeted by initiators most consistently develop a greater number of out-connections (A) and in-connections (B). **C & D.** Proportion of nodes with zero out-links (C) and in-links (D) connectivity among those targeted by initiators (left panel) and those not targeted (right panel). Results are shown as a function of the probability of proximity-based rewiring and retinal wave-based rewiring. The remaining probability (i.e. when the sum of the probabilities for proximity-based and retinal wave-based rewiring does not reach 1) corresponds to adaptive rewiring based on graph diffusion. For each combination of rewiring rule probabilities (e.g.: *p*_*proximity*_ = 0.5, *p*_*wave*_ = 0.2, *p*_*adaptive*_ = 0.3), results are obtained by averaging over 150 networks that evolve from 150 initially random networks.

In-link connectivity is generally larger for targeted nodes than for non-targeted ones. The difference is more pronounced at the lower frequencies of retinal wave occurrence. The increase in in-link connectivity is matched with a decrease in the out-connectivity in those nodes (Figure 7A). Among the nodes targeted and not targeted by initiators, Figures 7C and D show, respectively, the proportion of nodes that have zero out-connections and zero in-connections. Self-evidently, none of the nodes targeted by initiators in Figure 7D have zero in-degree. However, a significant proportion of non-targeted nodes has zero in-connectivity. Importantly, Figure 7C shows that the proportion of nodes with zero out-link connectivity is larger among nodes targeted by initiators than among non-targeted ones.

The result suggests that the effects of retinal waves on the differentiation of retinal ganglion also affects their direct neighbours. In the retina, the ganglion cells form recurrent circuits with amacrine cells (Xu et al., 2016). Mature amacrine cells are inhibitory and most lack an axon (Masland, 2012). It is possible that the differentiation of their morphology and function are driven by overexpression of Ca2+ in these cells, as a result of receiving the retinal wave activity. Intracellular calcium waves regulate the rate of axon extension (Rosenberg & Spitzer, 2011) and, at least in embryonic spinal neurons, the differentiation of inhibitory and excitatory cells. This occurs in a homeostatic manner, i.e. when calcium activity is low, more neurons tend to become excitatory and when it is high, inhibitory (Borodinsky et al., 2004).

Finally, we considered whether nodes targeted by initiators were more likely to become convergent hubs than the rest of the non-targeted nodes. Figure 8A shows that the proportion of convergent hubs among targeted nodes is larger than among nodes not targeted by initiators. Furthermore, Figure 8B shows the same for the proportion of convergent hubs within convergent-divergent units. The result suggests that the divergence created by retinal wave activity may prepare the ground for convergence at the level of their axon terminals in LGN (e.g. Hamos et al., 1987; Tavazoie & Reid, 2001) and beyond (Ringach, 2021), i.e. RGCs (divergent hubs in a convergent-divergent unit) preferably broadcast their input to convergent hubs in downstream areas (LGN and beyond), thereby reinforcing their role.

**Figure 8.**
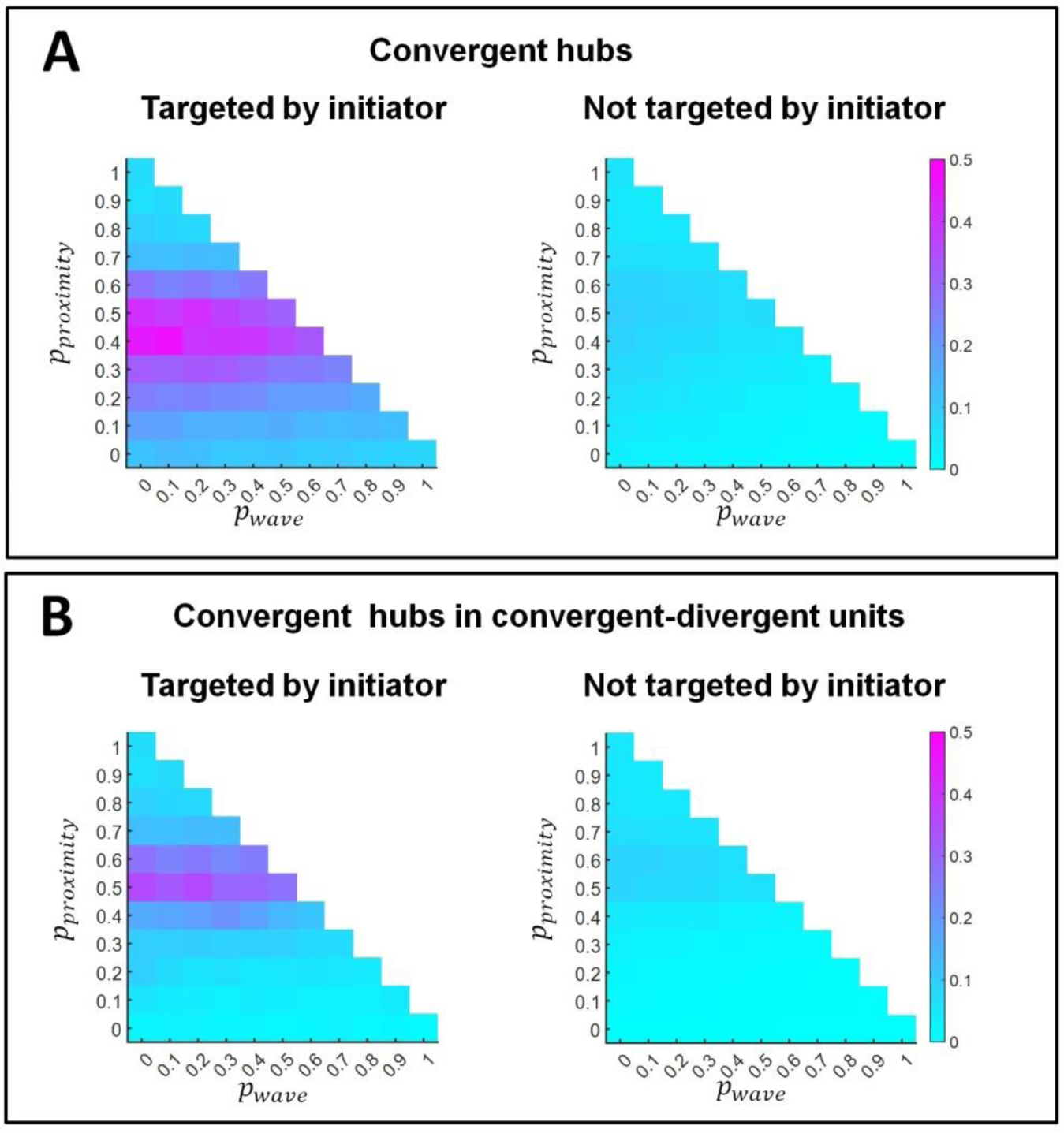
Proportion of convergent hubs (A), and convergent hubs within convergent-divergent units (B) among nodes targeted by the outlinks of retinal wave initiator nodes (left panels) and the rest of non-targeted nodes (right panels). Results are shown as a function of the probability of proximity-based rewiring and retinal wave-based rewiring. The remaining probability (i.e. when the sum of the probabilities for proximity-based and retinal wave-based rewiring does not reach 1) corresponds to adaptive rewiring based on graph diffusion. For each combination of rewiring rule probabilities (e.g.: *p*_*proximity*_ = 0.5, *p*_*w*__*ave*_ = 0.2, *p*_*adaptive*_ = 0.3), results are obtained by averaging over 150 networks that evolve from 150 initially random networks.

## Discussion

Applying a mixture of adaptive and spatial rewiring rules in neural networks with weighted and directed connections results in modular small-world networks with rich club effect, and convergent-divergent units (Li et al., 2023). Like in the present work, Li’s (2023) model relies entirely on spontaneous activity. This activity triggers synaptic plasticity mechanisms that compare well to our adaptive rewiring rules, i.e. growth cone behaviour at the tip of the axon or dendrite is influenced by the influx of calcium following postsynaptic activation (Kater et al., 1988, Kater et al., 1989, Kater and Guthrie, 1989, Kater and Guthrie, 1990, Kater et al., 1990, Jourdain et al., 2003); while spatial rewiring may be thought of as modelling gap junctions.

However, Li’s (2023) model does not have the capacity for sensory processing, a feature for which input systems are required. Considering this limitation, we are interested in how adaptive rewiring can help build the relevant input structures, based on spontaneous activity in the system. We focused on retinal wave activity, in particular on its role in the development of retinal ganglion cells as divergent hubs for projecting retinal activity into the visual system.

Initiator nodes, designated as retinal ganglion cells in an initially random network, received a boost to their outgoing activity, representing the effect of retinal waves. As a result, a significant proportion of initiator nodes developed into divergent hubs. Imposing retinal waves on the system did not interfere with the formation of complex network structures, including the formation of convergent-divergent units (Rentzeperis et al., 2022; Li et al., 2023).

The frequency with which the waves occur controls parametrically the proportion of initiator nodes that develop into divergent hubs. However, the largest proportions occur, not with the maximally frequent occurrence of waves, but in an intermediate regime, in accordance with their intermittent occurrence in the developing brain. The reason is that, while initiator nodes increasingly developed outgoing connectivity with the frequency of wave occurrence, this went at the expense of their incoming connectivity. To qualify as divergent hubs, incoming connections are necessary. With very prominent wave activity, initiator nodes increasingly lost their incoming connectivity. This result may suggest new insights into a topic that has been a conundrum in the field of retinal development: a large proportion of all ganglion cells dies naturally during development (De Montigny et al., 2023). While it might be intuitive to think that cell death is caused by insufficient input, our results suggest that an overdose of wave activity causes potential ganglion cells to lose their input, and die as a result.

The mechanism that turns initiator nodes into divergent hubs reduces the outgoing connectivity of their direct neighbours. At the same time, these nodes gain incoming connectivity (Figures 3B and 7). This effect may reflect the differentiation of some of these nodes into amacrine cells, which have many incoming connections but no axon. More generally, nodes targeted by initiator nodes are more likely to become convergent hubs (Figure 8A) and feature as such in convergent-divergent units (Figure 8B). This suggests that the retinal wave mechanism that effectuates divergence in the ganglion cells is also responsible for convergence downstream, in LGN and visual cortex. The result suggests that the divergence created by retinal waves could be linked to the formation of convergence in LGN. This is noteworthy, as convergence in LGN (Hamos et al., 1987; Tavazoie & Reid, 2001) and beyond (Ringach, 2021) is well-established empirically, and is held responsible for the diversity in receptive fields. Its origin has not been well-understood. It may be considered a serendipitous effect of our modelling strategy. The current result is limited in several ways. It abstracts from the differentiated ways remodelling of synapses in the developing brain takes place. It reduces the multistage complexity of retinal wave activity (Maccione et al, 2014) to a single process targeting initiator nodes only. By abstracting from this complexity, we aimed at the discovery of principles, which may play a basic role in more realistic future models.

Another limitation of the model is that it ignores the 3D spatial context and the impact of guidance cues on the navigation of axonal growth cones in network formation. Previous models (Calvo Tapia et al., 2020 and Li et al. 2023) offered a complementary third rewiring rule, termed “alignment” (Li et al., 2023) to represent these effects. The “alignment” rule allowed nodes, inter alia, to form chains. The rule was omitted here since it had little impact on the network structure (as opposed to its morphology). In future developments, our model could be combined with the simulation of axonal guidance and diffusive substances in physical space (Bauer et al., 2014). The timing of tectal and thalamic innervation and synapse formation significantly overlaps with the presence of retinal waves (Harada et al., 2007; Lalitha et al., 2020). Our results highlight the possibility that retinal waves promote axonal projections, while relying on the guidance cues to bring them to their destinations, LGN, pretectal nuclei, and superior colliculus. Similar mechanisms in these cell bodies might then arrange for further transport of retinal wave activity.

We conclude that by modelling retinal waves, we have achieved a systematic method to stochastically control the formation of outgoing hubs, or ganglia, in subgraphs of evolving complex networks. Our results suggest to that this very basic principle, which drives the formation of divergence in retinal ganglion cells, may also be responsible for the differentiation of amacrine cells and for convergence downstream the visual system. Ultimately, the retinal waves enable divergence and convergence to arise within the developing visual system.

## APPENDIX A

### Derivation of the consensus and advection kernels

In Rentzeperis et al. (2022), the authors extended the concept of diffusion in undirected graphs as presented in Jarman et al. (2017) to directed graphs that incorporated the principles of consensus and advection. Like diffusion, consensus (Ren et al., 2007) and advection (Chapman, 2015) involve the convergence of node values, called concentrations, towards a global state based on the local state of each node. For consensus, the rate of concentration change for node *i* is described as:

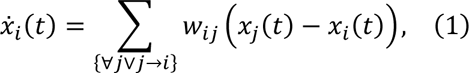

We define the in-degree Laplacian matrix, *L*_*in*_ = {*l*_(*ij*)_}, of the graph *G* as:

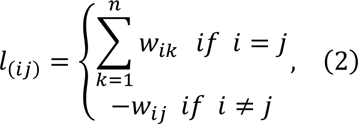

Substituting Equation 2 into Equation 1, the consensus dynamics in matrix form has the form:

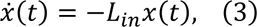

At time *t*, the nodes’ concentration is given by:

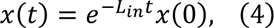

The advection dynamics governing node *i* can be mathematically expressed as follows:

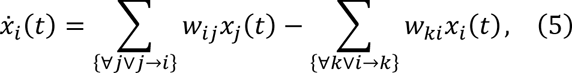

We define the out-degree Laplacian matrix, *L*_*out*_ = {*l*_*out*(*ij*)_}, of the graph *G* as:

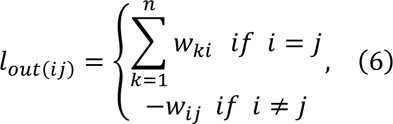

For advection, the rate of concentration change for node *i* is described as:

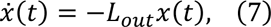

where the solution is:

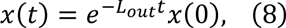

To establish the initial state of each node, we assign a concentration value of one to a particular node, and set the concentrations of the remaining nodes to zero. We then construct an identity matrix that represents these initial conditions. If we apply them to Equations 4 and 8, we obtain:

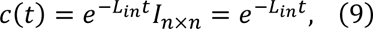

and:

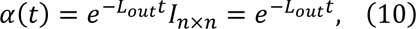

*c*(*t*) and *a*(*t*) are the consensus and advection kernels respectively, which represent communication intensity between nodes. These equations depend on graph laplacians, which change during network evolution due to network rewiring (i.e. laplacian matrices change as network topologies change due to rewiring). However, in our formulas we consider the graph laplacians to be constant (i.e., between rewiring iteration steps), for the sake of simplicity.

## References

Ackman, J. B., Burbridge, T. J., & Crair, M. C. (2012). Retinal waves coordinate patterned activity throughout the developing visual system. Nature, 490(7419), 219–225. 10.1038/nature11529

Adesnik, H., Bruns, W., Taniguchi, H., Huang, Z. J., & Scanziani, M. (2012). A neural circuit for spatial summation in visual cortex. Nature, 490(7419), 226–231. 10.1038/nature11526

Alexander, D. M., Trengove, C., Sheridan, P., & van Leeuwen, C. (2011). Generalization of learning by synchronous waves: from perceptual organization to invariant organization. Cognitive Neurodynamics, 5, 113–132. 10.1007/s11571-010-9142-9

Antonello, P. C., Varley, T. F., Beggs, J., Porcionatto, M., Sporns, O., & Faber, J. (2022). Self-organization of in vitro neuronal assemblies drives to complex network topology. Elife, 11, e74921. 10.7554/elife.74921

Arenas, A., Duch, J., Fernández, A., & Gómez, S. (2007). Size reduction of complex networks preserving modularity. New Journal of Physics, 9(6), 176–176. 10.1088/1367-2630/9/6/176

Arroyo, D. A., & Feller, M. B. (2016). Spatiotemporal features of retinal waves instruct the wiring of the visual circuitry. Frontiers in Neural Circuits, 10. 10.3389/fncir.2016.00054.

Bauer, R., Zubler, F., Hauri, A., Muir, D. R., & Douglas, R. J. (2014). Developmental origin of patchy axonal connectivity in the neocortex: a computational model. Cerebral Cortex, 24(2), 487–500. 10.1093/cercor/bhs327

Bender, E. A., & Williamson, S. G. (2010). Lists, decisions and graphs. S. Gill Williamson.

Blankenship, A.G., & Feller, M.B. (2010). Mechanisms underlying spontaneous patterned activity in developing neural circuits. Nature Reviews Neuroscience, 11, 18–29. 10.1038/nrn2759

Borodinsky, L. N., Root, C. M., Cronin, J. A., Sann, S. B., Gu, X., & Spitzer, N. C. (2004). Activity-dependent homeostatic specification of transmitter expression in embryonic neurons. Nature, 429(6991):523–30. doi: 10.1038/nature02518. PMID: 15175743. 10.1038/nature02518

Bullmore, E. & Sporns, O. (2009). Complex brain networks: graph theoretical analysis of structural and functional systems. Nature Reviews Neuroscience, 10, 186–198. 10.1038/nrn2575

Calvo Tapia, C., Makarov, V., & van Leeuwen, C. (2020). Basic principles drive self-organization of brain-like connectivity structure. Communications in Nonlinear Science and Numerical Simulation, 82, 105065. 10.1016/j.cnsns.2019.105065

Chapman, A. (2015). Advection on graphs. In A. Chapman (Ed.), Semi-Autonomous Networks: Effective Control of Networked Systems through Protocols, Design, and Modeling (pp. 3–16). Springer International Publishing. 10.1007/978-3-319-15010-9_1

de Montigny, J., Sernagor, E., & Bauer, R. (2023). Retinal self-organization: a model of retinal ganglion cells and starburst amacrine cells mosaic formation. Open Biology, 13(4), 220217. 10.1098/rsob.220217

D’Souza, S. & Lang, R.A. (2020).Retinal ganglion cell interactions shape the developing mammalian visual system. Development, 147(23). 10.1242/dev.196535

Galli L. & Maffei, L. (1988). Spontaneous impulse activity of rat retinal ganglion cells in prenatal life. Science, 242(4875), 90–91. 10.1126/science.3175637

Gong, P. & van Leeuwen, C. (2003). Emergence of scale-free network with chaotic units. *Physica A*, Statistical mechanics and its applications, 321, 679–688. 10.1016/s0378-4371(02)01735-1

Gong, P. & van Leeuwen, C. (2004). Evolution to a small-world network with chaotic units. Europhysics Letters, 67, 328–333. 10.1209/epl/i2003-10287-7

Haqiqatkhah, M.H. & van Leeuwen, C. (2022). Adaptive rewiring in non-uniform coupled oscillators. Network Neuroscience, 6 (1), 90–117. 10.1162/netn_a_00211

Hamos, J.E., Van Horn, S.C., Raczkowski, D., & Sherman, S.M. (1987). Synaptic circuits involving an individual retinogeniculate axon in the cat. Journal of Comparative Neurology, 259, 165–192. 10.1002/cne.902590202

Harada, T., Harada, C., & Parada, L. F. (2007). Molecular regulation of visual system development: more than meets the eye. Genes & Development, 21(4), 367–378. 10.1101/gad.1504307

Hellrigel, S., Jarman, N., & van Leeuwen (2019). Adaptive rewiring of weighted networks. Cognitive Systems Research, 55, 205–218. 10.1016/j.cogsys.2019.02.004

Hettiarachchi, N.T., Dallas, M.L., Pearson, H.A., Bruce, G., Deuchars, S., Boyle, J.P, & Peers, C. (2010). Gap junction-mediated spontaneous Ca^2+^ waves in differentiated cholinergic SN56 cells. Biochemical and Biophysical Research Communications, 397, 564–568. 10.1016/j.bbrc.2010.05.159

Himmelberg M.M., Tünçok E, Gomez J, Grill-Spector K, Carrasco M, & Winawer J. (2023)., Comparing retinotopic maps of children and adults reveals a late-stage change in how V1 samples the visual field. Nature Communications, 14(1), 1561. 10.1038/s41467-023-37280-8

Hoon, M., Okawa, H., Della Santina, L., & Wong, R. O. (2014). Functional architecture of the retina: development and disease. Progress in Retinal and Eye Research, 42, 44–84. 10.1016/j.preteyeres.2014.06.003

Jarman, N., Steur, E., Trengove, C., Tyukin, I., & van Leeuwen, C. (2017). Self-organisation of small-world networks by adaptive rewiring in response to graph diffusion. Scientific Reports, 7, 13518. 10.1038/s41598-017-12589-9

Jarman, N., Trengove, C., Steur, E., Tyukin, I., & van Leeuwen, C. (2014). Spatially constrained adaptive rewiring in cortical networks creates spatially modular small-world architectures. Cognitive Neurodynamics, 8, 479-497. 10.1007/s11571-014-9288-y

Jourdain, P., Fukunaga, K., & Muller, D. (2003). Calcium/calmodulin-dependent protein kinase II contributes to activity-dependent filopodia growth and spine formation. Journal of Neuroscience, 23(33), 10645–10649. 10.1523/jneurosci.23-33-10645.2003

Kater, S. B., Mattson, M. P., Cohan, C., & Connor, J. (1988). Calcium regulation of the neuronal growth cone. Trends in neurosciences, 11(7), 315–321. 10.1016/0166-2236(88)90094-x

Kater, S. B., Mattson, M. P., & Guthrie, P. B. (1989). Calcium-induced neuronal degeneration: a normal growth cone regulating signal gone awry (?). Annals of the New York Academy of Sciences, 568, 252–261. 10.1111/j.1749-6632.1989.tb12514.x

Kater, S. B., & Guthrie, P. B. (1989). The neuronal growth cone: calcium regulation of a presecretory structure. Society of General Physiologists series, 44, 111–122.

Kater, S. B., & Guthrie, P. B. (1990, January). Neuronal growth cone as an integrator of complex environmental information. In Cold Spring Harbor Symposia on Quantitative Biology (Vol. 55, pp. 359-370). Cold Spring Harbor Laboratory Press. 10.1101/sqb.1990.055.01.037

Kater, S. B., Guthrie, P. B., & Mills, L. R. (1990). Integration by the neuronal growth cone: a continuum from neuroplasticity to neuropathology. Progress in brain research, 86, 117–128. 10.1016/S0079-6123(08)63171-4

Katz, L. C., & Shatz, C. J. (1996). Synaptic activity and the construction of cortical circuits. Science, 274(5290), 1133–1138. 10.1126/science.274.5290.1133

Kerschensteiner, D. (2022). Feature detection by retinal ganglion cells. Annual Review of Vision Science, 8, 135–169. 10.1146/annurev-vision-100419-112009

Kumar, A., Rotter, S. & Aertsen, A. (2010). Spiking activity propagation in neuronal networks: reconciling different perspectives on neural coding. Nature Reviews Neuroscience, 11, 615–627. 10.1038/nrn2886

Kwok, H. F., Jurica, P., Raffone, A., & van Leeuwen, C. (2007). Robust emergence of small-world structure in networks of spiking neurons. Cognitive Neurodynamics, 1, 39–51. 10.1007/s11571-006-9006-5

Lalitha, S., Basu, B., Surya, S., Meera, V., Riya, P. A., Parvathy, S., Das, A. V., Sivakumar, K. C., Nelson-Sathi, S., & James, J. (2020). Pax6 modulates intra-retinal axon guidance and fasciculation of retinal ganglion cells during retinogenesis. Scientific Reports, 10(1), 16075. 10.1038/s41598-020-72828-4

Latora, V., & Marchiori, M. (2001). Efficient behavior of small-world networks. Physical review letters, 87(19), 198701. 10.1103/physrevlett.87.198701

Leicht, E. A., & Newman, M. E. (2008). Community structure in directed networks. Physical Review Letters, 100(11), 118703. 10.1103/physrevlett.100.118703

Leybaert, L., & Sanderson, M. J. (2012). Intercellular Ca^2+^ waves: mechanisms and function. Physiological Reviews, 92(3), 1359–1392. 10.1152/physrev.00029.2011

Li, J., Rentzeperis, I. & van Leeuwen, C. (2023). Functional and spatial rewiring jointly generate convergent-divergent units in self-organizing networks. PloS Computational Biology, 19(8), e1011325. 10.1371/journal.pcbi.1011325.

Luhmann, H. J. (2022). Neurophysiology of the developing cerebral cortex: What we have learned and what we need to know. Frontiers in Cellular Neuroscience, 3(15) 814012. 10.3389/fncel.2021.814012.

Maccione, A., Hennig, M. H., Gandolfo, M., Muthmann, O., van Coppenhagen, J., Eglen, S. J., Berdondini, L., & Sernagor, E. (2014). Following the ontogeny of retinal waves: pan-retinal recordings of population dynamics in the neonatal mouse. Journal of Physiology, 592(7), 1545–1563. 10.1113/jphysiol.2013.262840.

Masland R. H. (2012). The neuronal organization of the retina. Neuron, 76, 266–280. 10.1016/j.neuron.2012.10.002

Niell, C. M. & Scanziani, M. (2021). How cortical circuits implement cortical computations: Mouse visual cortex as a model. Annual Review of Neuroscience, 44, 517–546. 10.1146/annurev-neuro-102320-085825

Opsahl, T., Agneessens, F., & Skvoretz, J. (2010). Node centrality in weighted networks: Generalizing degree and shortest paths. Social networks, 32(3), 245–251. 10.1016/j.socnet.2010.03.006

Purves, D., Augustine, G., Fitzpatrick, D., Katz, L., LaMantia, A., McNamara, J., & Williams, S. (2001). Neuroscience 2nd edition. Sunderland, USA: Sinauer Associates.

Ren, W., Beard, R. W., & Atkins, E. M. (2007). Information consensus in multivehicle cooperative control. IEEE Control Systems Magazine, 27(2), 71–82. 10.1109/mcs.2007.338264

Rentzeperis, I., & Van Leeuwen, C. (2021). Adaptive rewiring in weighted networks shows specificity, robustness, and flexibility. Frontiers in Systems Neuroscience, 15, 580569. 10.3389/fnsys.2021.580569

Rentzeperis, I., Laquitaine, S. & van Leeuwen, C. (2022). Adaptive rewiring of random neural networks generates convergent-divergent units. Communications in Nonlinear Science and Numerical Simulation, 107, 106135. 10.1016/j.cnsns.2021.106135

Ringach, D. L. (2021). Sparse thalamocortical convergence. Current Biology, 31(10). 10.1016/j.cub.2021.02.032

Rosenberg, S. S., & Spitzer, N. C. (2011). Calcium signaling in neuronal development. Cold Spring Harbor Perspectives in Biology, 3(10), a004259. 10.1101/cshperspect.a004259

Shatz, C. J. (1996). Emergence of order in visual system development. Journal of Physiology-Paris, 90(3-4), 141–150. 10.1016/s0928-4257(97)81413-1

Shaw, G. L., Harth, E. & Scheibel, A. B. (1982). Cooperativity in brain function: assemblies of approximately 30 neurons. Experimental Neurology, 77, 324–358. 10.1016/0014-4886(82)90249-7

Song Y.H., Hwang, Y.S., Kim, K., Lee, H.R., Kim, J.H., Maclachlan, C., Dubois, A., Jung, M.W., Petersen, C.C.H., Knott, G., Lee, Suk-Ho, & Lee, Seung-Hee (2020). Somatostatin enhances visual processing and perception by suppressing excitatory inputs to parvalbumin-positive interneurons in V1. Science Advances, 6(17), eaaz0517. doi: 10.1126/sciadv.aaz0517.

Tavazoie, S.F., & Reid, R.C. (2000) Diverse receptive fields in the lateral geniculate nucleus during thalamocortical development. Nature Neuroscience, 3(6), 608–616. 10.1038/75786. PMID: 10816318.

van den Berg, D., Gong, P., Breakspear, M., & van Leeuwen, C. (2012). Fragmentation: Loss of global coherence or breakdown of modularity in functional brain architecture? Frontiers in Systems Neuroscience, 6, 20. 10.3389/fnsys.2012.00020

van den Berg, D. & van Leeuwen, C. (2004). Adaptive rewiring in chaotic networks renders small-world connectivity with consistent clusters. Europhysics Letters, 65, 459–464. 10.1209/epl/i2003-10116-1

van den Heuvel, M. P., & Sporns, O. (2011). Rich-club organization of the human connectome. Journal of Neuroscience, 31, 15775–15786. 10.1523/jneurosci.3539-11.2011

Weiss, S., Clamon, L.C., Manoim, J.E., Ormerod, K.G., Parnas, M., & Littleton, J.T. (2022). Glial ER and GAP junction mediated Ca^2+^ waves are crucial to maintain normal brain excitability. Glia, 70(1), 123–144. 10.1002/glia.24092.

Xu, H.P., Burbridge, T.J., Ye, M., Chen, M, Ge, X., Zhou, Z.J., & Crair, M.C. (2016). Retinal wave patterns are governed by mutual excitation among starburst amacrine cells and drive the refinement and maintenance of visual circuits. Journal of Neuroscience, 36(13), 3871–3886. 10.1523/jneurosci.3549-15.2016

Zamora-López, G., Zhou, C., and Kurths, J. (2010). Cortical hubs form a module for multisensory integration on top of the hierarchy of cortical networks. Frontiers in Neuroinformatics, 4, 1. 10.3389/neuro.11.001.2010

